# Multiscale Mechanical Characterization and Computational Modeling of Fibrin Gels

**DOI:** 10.1101/2022.12.06.519227

**Authors:** Julian M. Jimenez, Tyler Tuttle, Yifan Guo, Dalton Miles, Adrian Buganza-Tepole, Sarah Calve

## Abstract

Fibrin is a naturally occurring protein network that forms a temporary structure to enable remodeling during wound healing. It is also a common tissue engineering scaffold because the structural properties can be controlled. However, to fully characterize the wound healing process and improve the design of regenerative scaffolds, understanding fibrin mechanics at multiple scales is necessary. Here, we present a strategy to quantify both the macroscale (1 – 10 mm) stress-strain response and the deformation of the mesoscale (10 – 1000 μm) network structure during unidirectional tensile tests. The experimental data is then used to inform a computational model to accurately capture the mechanical response of fibrin gels. Simultaneous mechanical testing and confocal microscopy imaging of fluorophore-conjugated fibrin gels revealed up to an 88% decrease in volume coupled with increase in volume fraction in deformed gels, and non-affine fiber alignment in the direction of deformation. Combination of the computational model with finite element analysis enabled us to predict the strain fields that were observed experimentally within heterogenous fibrin gels with spatial variations in material properties. These strategies can be expanded to characterize and predict the macroscale mechanics and mesoscale network organization of other heterogeneous biological tissues and matrices.

**Statement of Significance:** Fibrin is a naturally-occurring scaffold that supports cellular growth and assembly of *de novo* tissue and has tunable material properties. Characterization of meso- and macro-scale mechanics of fibrin gel networks can advance understanding of the wound healing process and impact future tissue engineering approaches. Using structural and mechanical characteristics of fibrin gels, a theoretical and computational model that can predict multiscale fibrin network mechanics was developed. These data and model can be used to design gels with tunable properties.

## 1 Introduction

Fibrin is an insoluble protein network that serves critical roles in thrombosis and wound healing. During dermal wound healing, fibrinogen polymerizes into a fibrin network that creates a temporary structure to promote hemostasis, and provides an environment that mediates cellular adhesion, migration, proliferation, and differentiation, which is crucial for the initiation of extracellular matrix (ECM) remodeling [1]. Because dermal wounds are frequently subjected to large deformations during healing, it is important to understand how fibrin mechanics contributes to the mechanobiology of the dermal wound healing process. Additionally, fibrin is widely implemented in tissue engineering and cell culture applications as it can be polymerized into gels with controlled structural properties (*e.g*. fiber length, diameter, fibrin volume fraction) by modulating factors such as fibrinogen, thrombin and Ca^2+^ concentration, ionic strength, pH, and temperature [1,2][3], generating a range of material properties.

The mechanics of fibrin have been studied at various hierarchical scales, from the molecular interactions between fibrin monomers, to the mesoscale (10 – 1000 μm) fiber network features [4,5], to the macroscale (1 – 10 mm) stress-strain response of whole gels [4]. Atomic force microscopy (AFM) analysis of individual fibrin fibers measured moduli ranging from 1.0-9.7 MPa [6–9], which decreased with increasing fiber diameter [7]. In contrast, the moduli of fibrin networks measured via AFM was lower than individual fibers, and depended on tip geometry, ranging from 1-12 kPa for microscale probes and 10-1000 kPa for nanoscale probes [10]. As a bulk material, fibrin gels behave viscoelastically when investigated via rheology and tensile testing [10–19]. The storage modulus was shown to be much greater than the loss modulus [10,12,13,15–19], demonstrating a mostly elastic mechanical response, as well as strain stiffening and strain-enhanced stress relaxation [14].

The organization of the fibrin network changes in response to loading. At the mesoscale, the orientation distribution of fibrin fibers polymerized in static conditions, *i.e*. no applied tensile or compressive loading, is isotropic [2,20–22]. In response to shear and tensile tests, the fibrin network aligns in the direction of deformation and fibers becomes more densely packed [4]. Deformed fibrin matrices exhibit volume loss [4], and fiber alignment in the direction of deformation has been shown to be non-affine [23]. It is clear that the mechanical environment influences fiber network organization, and vice versa; however, the simultaneous correlation between fiber orientation and network structure and the mechanical response at the meso- and macroscales have not been directly assessed.

Multiscale models have been used to describe and predict how macroscale deformation influences the mesoscale structures of hydrogel networks [24–26]. Conversely, multiscale models can help elucidate how structural changes at the mesoscale during deformation influence the macroscale tissue mechanics [4]. Computational models are also advantageous to leverage knowledge from idealized testing conditions in order to describe the mechanics of more complex and physiologically relevant systems, such as *in vivo* wound deformations following dermal injury, which are challenging to characterize empirically. Various models have been proposed to model the multiscale mechanical behavior of 3D hydrogel networks. For example, discrete fiber networks (DFN) models with defined parameters for fiber length, diameter, orientation distribution, and volume fraction within a representative volume element (RVE) have been proposed [27–29]. DFN models can then be used to inform analytical or data-driven models representative of an average RVE response [29]. Alternatively, fitting analytical models at the macroscale that also incorporate mesoscale information such as fiber orientation distribution and volume fraction is also possible [30]. To better inform the computational models, experimental characterization of mesoscale structural properties of fibrin gels under tensile deformation, alongside the macroscale stress and strain response, is needed.

Here, we present a coupling of the mesoscale fibrin network organization at varying strains with macroscale stress and strains of gels, which is used to inform a multiscale computational model. We demonstrate the fibrinogen concentration-dependent and viscoelastic mechanical behavior of these gels and the non-affine deformation of the fiber network. This is used to build a computational model based on structural characterization of fibrin networks under tension that enables us to accurately predict the strain response of heterogeneous fibrin gels composed of regions with different material properties. We anticipate this framework, which couples experimental and computational models, will facilitate a better understanding of multiscale mechanics of fibrin clots and can be adapted to other protein matrices and expanded to study the mechanobiology of dermal wound healing.

## 2 Materials and methods

### 2.1 Fibrin gel preparation

Homogeneous fibrin gels were prepared to final fibrinogen concentrations of 2 and 4 mg/mL by combining human fibrinogen (FIB3, Enzyme Research Laboratories) and Alexa Fluor (AF) 488 or 546 conjugated human fibrinogen (F13191, Molecular Probes) at a 1:10 fluorescent to non-fluorescent fibrinogen ratio, 1 μL 2M CaCl_2_, and 0.5 U thrombin (HT 1002A, Enzyme Research Laboratories) per mg fibrinogen in phosphate buffered saline (PBS). Immediately after mixing, the solution was pipetted into molds containing frames that enabled tensile testing (Fig. 1a-c) and allowed to polymerize for 5 minutes. Molds (2.5 × 6.6 × 0.75 mm) were designed in SolidWorks and 3D printed (Stratasys J750) with a photopolymer (VeroClear, Stratasys) that is similar to acrylic. Frames (6 × 12 mm) were drawn in Adobe Illustrator and laser cut from 100 μm thick polyethylene terephthalate using a Speedy 360 laser cutter (Trotec). Polyethylene (PE) blocks (coarse grade, Bel-Art Products) were manually cut to approximately 1.5 mm × 2.0 mm × 1.5 mm. To increase wettability of PE by the fibrinogen solution, the blocks were submerged in 70% ethanol for 5 minutes at room temperature (RT), submerged in 12.91 mg/mL fibrinogen solution for at least 1 hour, then glued to the laser-cut frames with cyanoacrylate adhesive.

**Figure 1.**
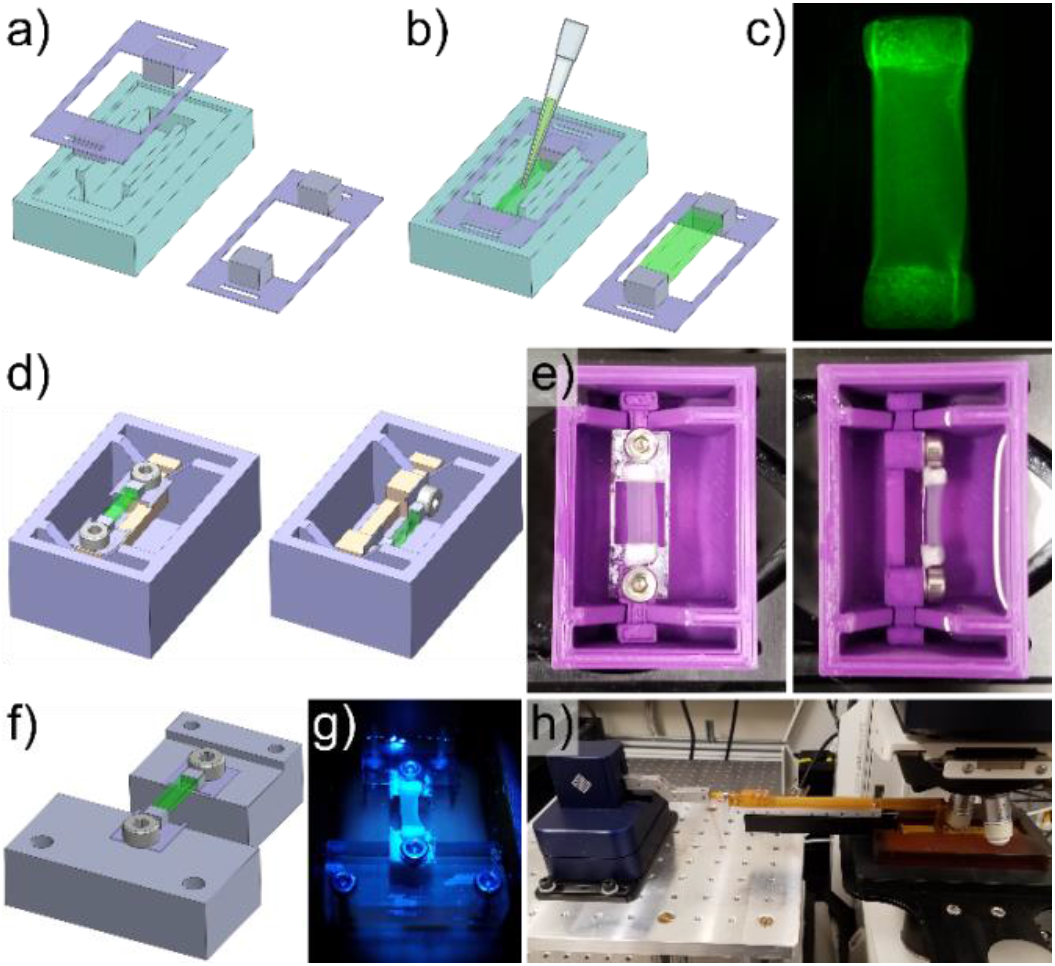
Fibrin gel polymerization and imaging setup. Porous polyethylene blocks were cut to size, coated with fibrinogen, and adhered to polyethylene terephthalate frames which were then loaded into molds (a). Immediately after combining thrombin and fibrinogen, the solution was pipetted into the frame-loaded molds prior to polymerization, and the frame-gel assembly was removed from the molds following fibrin polymerization (b). A representative 2 mg/mL fibrin gel with visible porous polyethylene blocks (c). A bath with a rotatable mounting system was designed (d) and 3D printed (e) to facilitate imaging of fibrin gels from 2 different orientations. A separate mounting system was designed (f), 3D printed (g), and combined with a micromanipulator (h) to image the fibrin gels during deformation while minimizing vibrations to improve image resolution.

### 2.2 Static, orthogonal fibrin gel imaging

To view the 3D mesoscale organization of the fibrin network, an orthogonal rotation bath (Fig. 1d,e) was used to image the gels in the *x* – *y* and *x* – *z* planes. This was necessary to accurately capture the 3D network fiber orientation distribution since resolution in the *z*-direction was lower than in the *x* – *y* plane with the available objectives. Image stacks were taken with a STELLARIS 5 confocal microscope (Leica), using an Apochromatic 63 × (NA = 0.90) water-immersible objective with a digital zoom of 6. Image stacks (29.4 × 29.4 × 29.4 μm) with resolutions of 1024 ×1024 × 70 px were generated at 0°and 90°rotation around the *x*-axis. Both 2 mg/mL (n=3) and 4 mg/mL (n=3) gels were imaged.

### 2.3 Stress-relaxation experiments

Two 50 × 50 μm fiducial markers were photobleached on the surface of the fibrin gels 200 μm apart, to a depth of 200 μm, using a STELLARIS confocal microscope and an Apochromatic 10× (NA = 0.40) water-immersible objective. Initial full-thickness fibrin gel cross-sections were imaged that encapsulated the entire region between the photobleached markers at 10×. Then fiber network image stacks were taken at 63 × following the procedure described in *Section 2.2*; however, only in the *x* – *y* configuration. Due to microscale vibrations that interfered with the simultaneous recording of force and confocal images, two experiments were performed to: (1) observe strain-dependent changes in fibrin network organization and (2) measure stress relaxation of the fibrin gels. A FemtoTools micromanipulator system (FT-RS1002) was used to apply unidirectional tensile deformations of the fibrin gels in both experiments [31], using different mounting systems. A custom slider assembly (without a microforce sensor) was used for experiment (1) (Fig. 1f-h). A custom floating-platform mounting system with a microforce sensor was used for experiment (2) (Fig. 2c-d). Fibrin gels were deformed to displacements of 1, 2, and 3 mm at a rate of 66 μm/s (approximately 0.01 s^-1^), with relaxation times of 60 min for each step. Both 2 mg/mL (n=3) and 4 mg/mL (n=3) gels were tested for each experiment (n=12 fibrin gels total).

**Figure 2.**
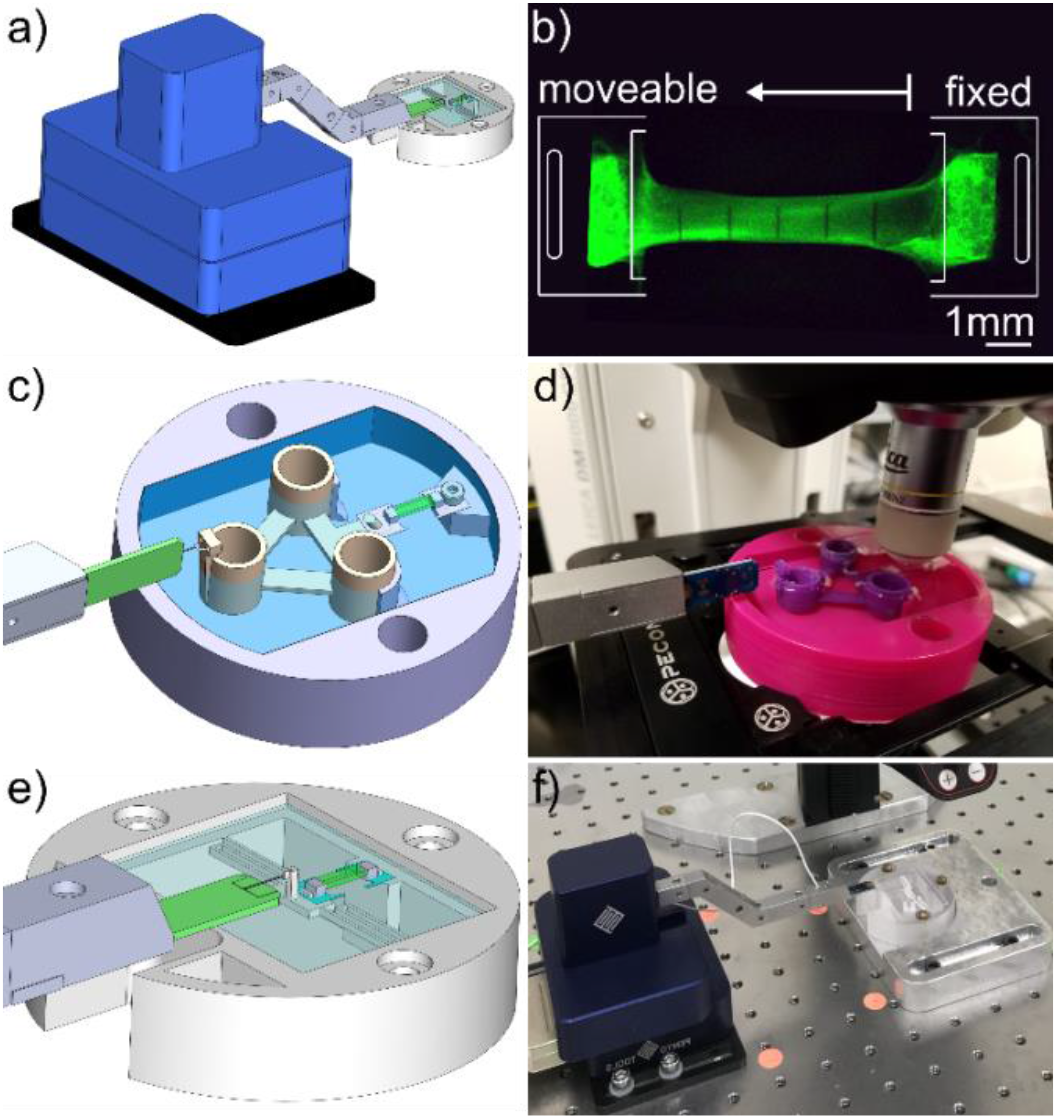
Fibrin gel mechanical testing setup. A micromanipulator and micro-force sensor were used to apply deformation and record force response from the gels (a). Fibrin gels were mounted with one fixed end and one moveable end, and gel mounting frame edges were removed prior to deformation (b). Two different mounting systems were used for mechanical testing. The time required for stress-relaxation tests (*Section 2.3*) necessitated a different mounting system that did not require baseline testing in addition to mounted sample testing. Therefore, a floating platform mounting system was designed (c) and used with the confocal microscope to image fibrin gels at different levels of deformation (d). For strain-rate dependent testing (*Section 2.4*) a spring mounting system described in [31] was designed (e) and used with deformation recorded with a dissecting microscope (f).

#### 2.3.1 Stretch-dependent matrix organization

A custom designed slider assembly and mounting system (Fig. 1f-h) was built to minimize microscale vibrations and enable confocal microscope imaging during sample deformation. For each level of deformation, full thickness image stacks were captured at 5, 30, and 50 minutes, at 10×, after individual displacements were applied to measure applied stretch and sample volume and cross-sectional area. Image stacks to capture mesoscale organization were also acquired at 15 and 40 minutes, using a 63× objective. Imaging was performed at more than one time point to check for time-dependent changes in matrix mesoscale and macroscale gel organization.

#### 2.3.2 Stress-relaxation tests

Mechanical testing was performed with the FemtoTools micromanipulator and 100 mN capacity microforce sensor probe (FT-S100000). A custom designed floating-platform mounting system was designed (Fig. 2c,d) to allow mechanical testing with minimal friction during deformation. Preliminary testing revealed that the force required to pull the unmounted floating platform was indistinguishable from sensor noise. Initial mesoscale fiber network and full thickness image stacks were taken prior to mechanical testing. At each level of deformation, full thickness image stacks were captured 50 minutes after displacements were applied.

### 2.4 Strain rate dependent mechanical testing

To measure macroscale stress-strain response, tensile tests were performed on 2 mg/mL fibrin gels at displacement rates of 6.6 (n=3), 66 (n=3), and 660 (n=4) μm/s (approximate strain rates of 0.001, 0.01, and 0.1 s^-1^) to a displacement of 6000 μm using the FemtoTools setup. Initial mesoscale fiber network and full thickness image stacks were obtained prior to mechanical testing. Five 41 × 533 μm fiducial vertical lines were photobleached on the surface of the fibrin gels 1000 μm apart, to a depth of 200 μm, using a STELLARIS confocal microscope and an Apochromatic 10× (NA = 0.40) water-immersible objective. The micromanipulator and force sensor described above were used with the mounting system in Fig. 2e-f [31]. Baseline tests, in which forces were measured with no sample mounted were performed after each test to subtract from the recorded mounted force data as previously described [31]. Deformation was recorded continuously throughout the tests with a INFINITY3-3URC camera (Lumenera) mounted on an M80 dissecting microscope (Leica Microsystems), at a frame rate of approximately 10 frames per second.

### 2.5 Heterogenous gel experiments

Heterogeneous constructs, in which a central inclusion of fibrin was surrounded by a different concentration of fibrin, were made using 2 mg/mL AF488-labeled fibrin, and 4 mg/mL fibrin with equal parts AF488 and AF546-labeled fibrin. First, the outer material was pipetted into a 4 mm × 6.6 mm × 0.75 mm mold, with inclusion material pipetted into the center of the gel 10s later. A 2 mg/mL gel with 4 mg/mL inclusion (n=1), and 4 mg/mL gel with 2 mg/mL inclusion (n=1) were generated. A 5 × 5 grid of 100 × 100 μm squares was photobleached 250 μm deep into the surface of the gels to encapsulate the inclusion and surrounding area. Initial full thickness image stacks were taken prior to mechanical testing. Mechanical tests were performed using the floating platform (Fig. 2c-d). Deformation was applied at a rate of 66 μm/s to a displacement of 3000 μm. Deformation was visualized as described in *Section 2.4*.

### 2.6 Data analysis

#### 2.6.1 Image analysis

Confocal images were deconvoluted using the Leica LIGHTNING settings in LAS X Life Science Microscope Software optimized for high resolution. Full thickness image stacks were used to measure distance, cross-sectional area, and volume between the photobleached markers. To measure applied tensile stretch of individual samples, fiducial marker location was tracked in MATLAB (MathWorks) by mean marker pixel location. Cross-sectional area and volume were measured in ImageJ (NIH) by cropping the image between fiducial markers, interpolating voxel intensity in the *z*-direction to make an isovoxel image stack, re-slicing to view the *y* – *z* orientation, manual thresholding to create a binary image, then applying a series of dilation, close, fill holes, and erosion functions to isolate the sample from background. Cross-sectional area was calculated as the product of pixel area and number of segmented pixels for a given *x*-slice. Volume was calculated as product of voxel volume and the total number of segmented voxels between photobleached markers.

Volume fraction was calculated as the ratio of fibrin voxels to total image voxels after binarizing the image stacks, acquired as described in above, using the Auto Local Threshold function in ImageJ. 2D orientation distributions were determined for 500 × 500 × 70 px cropped regions of the mesoscale fiber network image stacks using the OrientationJ [32] plugin in ImageJ. Orientation distributions were fit with a probability density function defined as a weighted summation of isotropic and von Mises distributions (Eq.(1)) [33],

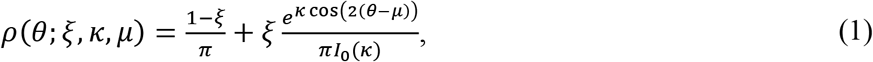

where ρ is the probability density function, θ is the fiber orientation (0 is in the *x*-direction), *ξ* is the weighting parameter, *κ* is the fiber alignment strength factor, μ is the preferred fiber orientation, and I_0_ is the modified Bessel function of the first kind of order zero. This distribution was fit to the RVE orientation distributions using `fminsearch` in MATLAB.

For gels imaged on the dissecting microscope (*Section 2.4* strain-rate study and *Section 2.5* heterogeneous gels), fiducial markers were tracked using the Manual Tracking plugin in ImageJ.

#### 2.6.2 Stress-strain response

The change in distance between fiducial markers measured in *Section 2.6.1*, was used to calculate stretch, λ, (length divided by initial length). Nominal stress, or first Piola–Kirchhoff stress (*P*_11_), was calculated by dividing force data by the initial cross-sectional area. Tangent moduli were fit to stress-stretch data (for stretches ranging from 1 to 1.4) using `regress` in MATLAB. Tangent moduli are used here as a method of comparison for experimental data and are not intended to be a constitutive description of the underlying material.

### 2.7 Constitutive modeling

#### 2.7.1 Nonlinear viscoelastic response of fiber networks

A hyper-viscoelastic model with two viscous branches motivated by the Holzapfel-Gasser-Ogden model [34] was proposed to model the fibrin gels following our previous work [30]. The strain energy density function of a fiber network of *n_f_* fibrin fibers with orientation *θ_i_* sampled from a weighted summation of isotropic and von Mises distributions in Δ*θ* intervals is captured by

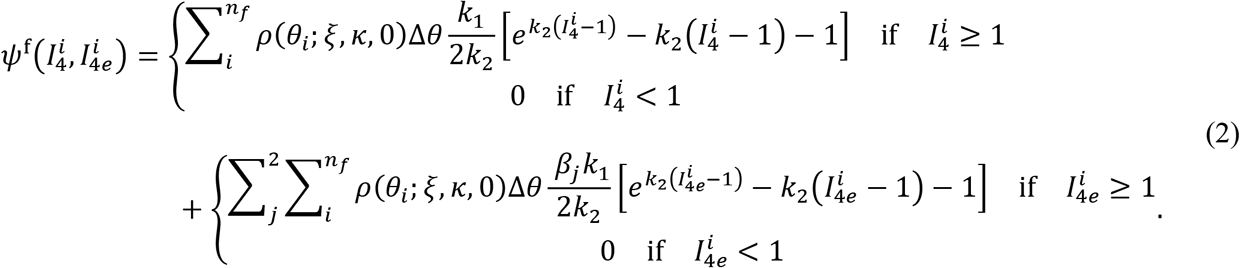

The stiffness, nonlinearity, and viscosity of the fibers are denoted by *k*_1_, *k*_2_ and *β_j_*, respectively. Each fibrin fiber contributes energy from an equilibrium elastic component (the first term in Eq.(2)) and a viscoelastic component (the second term in Eq.(2)). The equilibrium component of the *i^th^* fiber is expressed by a nonlinear function of the square of the total fiber stretch, 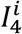, and the viscoelastic properties of that fiber are captured by another two nonlinear viscous branches depending on the elastic component of the fiber stretch 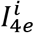. A relaxation time *τ_j_* is defined for the *j^th^* viscous branch. The update of the internal variables (viscous fiber stretches) assumes that the relaxation rate is proportional to the stress on that fiber [35], leading to

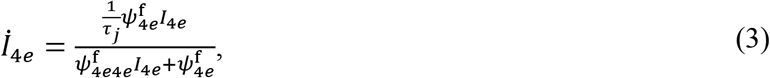

where

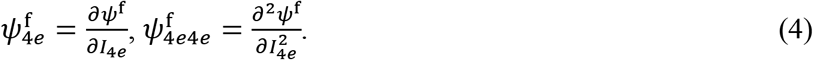

The viscous branches in Eq.(2) can thus be evolved in time by integrating Eq.(3).

The initial orientation distribution was assigned according to the experimental data. To capture the non-affine reorientation of fibers, a micromechanics approach is introduced in *Section 2.7.3*.

#### 2.7.2 Macroscale response of fibrin gels

The fibrin gels are regarded as a fiber-reinforced composite. Fibrin fibers are assumed to be embedded in an isotropic ‘ground substance’. The total strain energy density function includes the ground substance contribution, a volumetric energy term, and a weighted contribution from the fiber network taking into consideration the initial volume fraction, *ζ*,

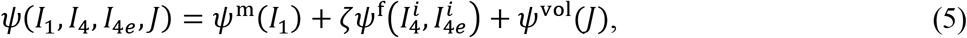

The isotropic component *ψ*^m^ is neo-Hookean parameterized by *k*_0_, and it is a function of the first invariant of the deformation *i.e*.

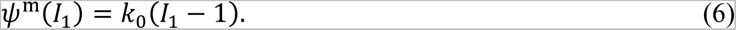

The volumetric term *ψ*^vol^ is

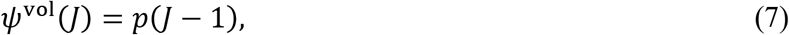

where *J* denotes the volume change and *p* can either come from the volumetric strain energy or be solved for from boundary conditions in the limit of incompressible behavior.

Calculation of the stress from the strain energy was done from standard arguments by differentiating the strain energy with respect to the corresponding strain measures. Explicit derivations are provided in the Supplement Eqs. (S1, S2).

#### 2.7.3 Non-affime reorientation of fibers

In fibrin gels, fibers in the network deformed non-affinely [23]. To capture this behavior, the micromechanics approach reported in [36] was adopted. Based on our observations and previous work on fibrin gels [25], fibers orient in the direction of the principal stretch direction *θ_c_* and this alignment increases as the maximum principal stretch *λ_c_* increases,

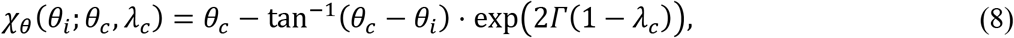

where *χ_θ_* denotes the mapping from the original orientation of the *i^th^* fiber (*θ_i_*) to its new reference orientation, with Γ the only parameter. Parameter Γ was optimized to replicate the changes in orientation distribution from the stretch-dependent matrix reorganization experiments.

#### 2.7.4 Constitutive model fit

To fit the strain energy parameters (*k*_0_, *k*_1_, *k*_2_, *β*_1_, *β*_2_, *τ*_1_, *τ*_2_), we first extracted the equilibrium response from the stress relaxation tests of fibrin gels and fit the subset of parameters (*k*_0_, *k*_1_, *k*_2_) for the 2mg/mL and 4mg/mL gels separately using the global optimizer `blackbox` in Python [37]. To balance between fitting each sample separately as opposed to finding the average response across all gels of a given fibrin concentration, individual parameters per sample were fit with the addition of the following correlation terms

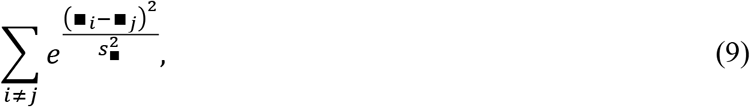

where ∎ indicates a given parameter: *k*_0_, *k*_1_, or *k*_2_, *β*_1_, *β*_2_, *τ*_1_ and *τ*_2_, were fit to match the transient portion of the stress relaxation test data. Similar to the equilibrium response, individual parameters were fit per sample with the addition of correlation terms across samples of a given fibrin concentration, analogous to Eq. (9).

### 2.8 Finite element analysis

A finite element code implemented in C++ and reported in [30] was used to simulate the deformation of fibrin gels in response to tensile testing. Mechanical equilibrium was reached at every time step by solving the stress divergence equation of **∇_*x*_** · ***σ*** = 0 plus boundary conditions. Specific simulations are described next.

A homogeneous and two heterogenous simulations were conducted. A 3.3 mm × 1.8 mm rectangle which represents a quarter of the whole gel (Fig. S3e) was generated for the homogeneous gels; a 6.6 mm × 3.2 mm rectangle with a 3.0 mm × 1.9 mm ellipse was used for the 2mg/mL fibrin gel with the 4 mg/mL inclusion; and a 6.8 mm × 3.8 mm rectangle with a 3.2 mm × 2.2 mm ellipse was used for the 4mg/mL fibrin gel with the 2 mg/mL inclusion. Structured quadrilateral meshes were generated by ABAQUS (Dassault Systemes Simulia Corp) for discretization. For the homogeneous gel, *x*- and *y*-symmetry boundary conditions were imposed on the left and bottom sides of the 3.3 mm × 1.8 mm domain. For heterogenous gels, clamped boundary conditions were enforced on the left boundary of the geometries, and the right boundary of all the geometries were displaced based on the displacement of fiducial points from experiments. The displacement of the fiducial points was used and not the clamp-to-clamp displacement since the latter was not an accurate representation of the stretch applied to the gels. The averaged fitting parameters of 2 mg/mL or 4 mg/mL fibrin gels (Fig. S3c) were used as material properties in the finite element simulations.

Two 3-step loading and stress relaxation tests were simulated for the homogeneous gels of 2 mg/mL and 4 mg/mL fibrin at strain rate of 0.01 s^-1^, analogous to the experiment. The stress near the bottom left corner of the quarter gel geometry, *i.e*. near the center of the homogeneous gel, was measured in order to compare the finite element results against the analytical model (Eq. (S1)). The heterogeneous geometries were stretched at a rate of 0.01 s^-1^ for 45 s and relaxed for 5 min to replicate the experimental conditions.

### 2.9 Statistical analysis

Comparison of orientation distribution parameters fit to Eq. (1) from the rotated fibrin gel image stacks (*Section 2.2*) was performed via 2-way ANOVA, with fibrinogen concentration and imaging plane (within-subjects) as variables. Comparison of stretch-dependent matrix organization parameters (*i.e*. volume fraction and orientation distribution parameters) was performed via 2-way ANOVA, with fibrinogen concentration and displacement (within-subjects) as sources of variation. Comparison of peak and relaxed moduli from stress-relaxation tests was performed with two-sample t-tests. Comparison of moduli from strain-rate dependent mechanical testing was performed with a 1-way ANOVA, with strain rate as the source of variation. Multiple comparison tests were conducted using of Tukey-Kramer post-hoc analysis. Statistical analyses were performed in MATLAB using `anovan` for n-way ANOVAs, `ranova` for n-way ANOVAs containing a within-subject source of variation, `ttest2` for two-sample t-tests, and `multcompare` for post-hoc multiple comparisons.

## 3 Results

### 3.1 Static fibril organization

To view the full 3D orientation distribution of fibrin fibers, and assess the validity of assuming radial symmetry of fiber orientation distribution around the *x*-axis, orthogonal orientations of the gels were imaged as described in *Section 2.2*. No noticeable differences in matrix organization were seen between image stacks obtained in the *x* – *y* and *x* – *z* orientations (Fig. 3a-d, representative slices). Mean fiber orientation distributions showed no discernible trends in anisotropy based on imaging plane (Fig. 3e,f). Eq. (1) was fit to the orientation distributions (Fig. 3g,h, representative fits) to uncover quantitative differences in fibrin orientation based on concentration or imaging plane. While there appeared to be a preferential orientation in the *x*-direction, there was no significant difference between fibrinogen concentration or imaging plane for any distribution parameters.

**Figure 3.**
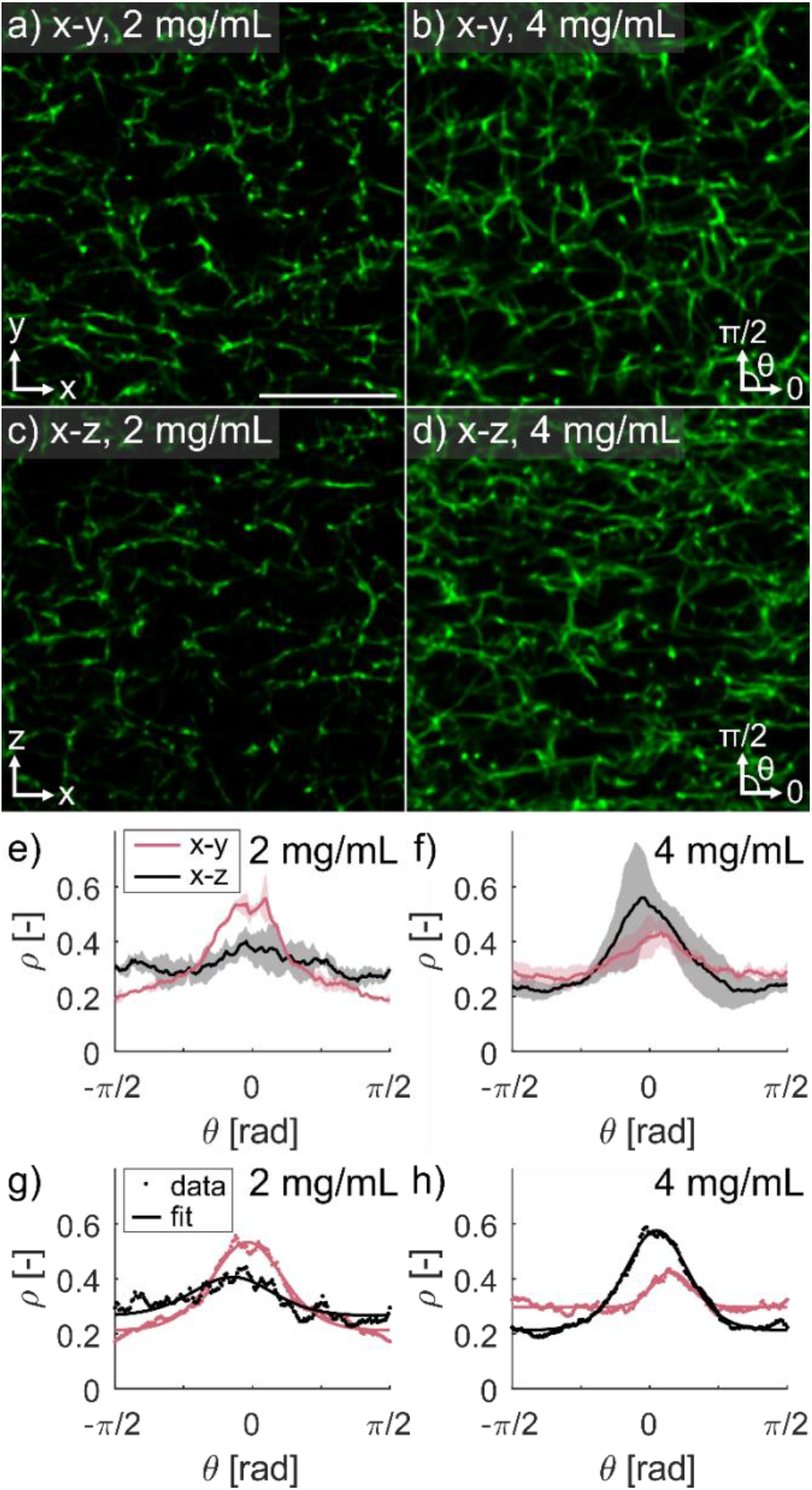
Undeformed fibrin gels did not show significant differences in 2D fiber orientation distributions based on imaging plane. Meso-scale imaging of fibrin gels was performed using the mounting setup described in Figure 1d-e. Representative image slices of the *x* – *y* plane of 2 mg/mL (a) and 4 mg/mL (b) gels, and representative image slices of the *x* – *z* plane of 2 mg/mL (a) and 4 mg/mL (b) gels (scale bar = 10 μm). Mean +/− standard deviation of orientation distributions of the *x* – *y* and *x* – *z* planes for 2 mg/mL (e) and 4 mg/mL (f) gels, with representative orientation distribution fits for a 2 mg/mL (g) and 4 mg/mL (h) gel. Greater alignment in the *x*-direction is seen in the *x* – *y* plane for 2 mg/mL gels but not seen in the *x* – *z* plane for 4 mg/mL gels. Statistical analysis of orientation distribution fit parameters revealed no differences for any parameters based on imaging plane or gel concentration.

### 3.2 Deformation-dependent fibril organization

Gels were deformed to quantify how the organization of fibrin fibers change as a function of displacement and fibrinogen concentration. Fibrin volume fraction, *ζ*, was higher in gels made with 4 mg/mL fibrinogen than 2 mg/mL gels, and matrix density visibly increased with increasing displacement (Fig. 4a-d). Volume fraction increased significantly with both displacement (p=1.3E-9) and fibrinogen concentration (p=1.4E-3) (Fig. 4e). Fibers aligned with the direction of unidirectional deformation (Fig. 4a-d), which is supported by the increase in prominence of the orientation distributions peaks in the direction of deformation (Fig. 4f,g). Orientation distribution fits show that the distribution function accurately captured the orientation distribution shape at all levels of displacement (Fig. 4h).

**Figure 4.**
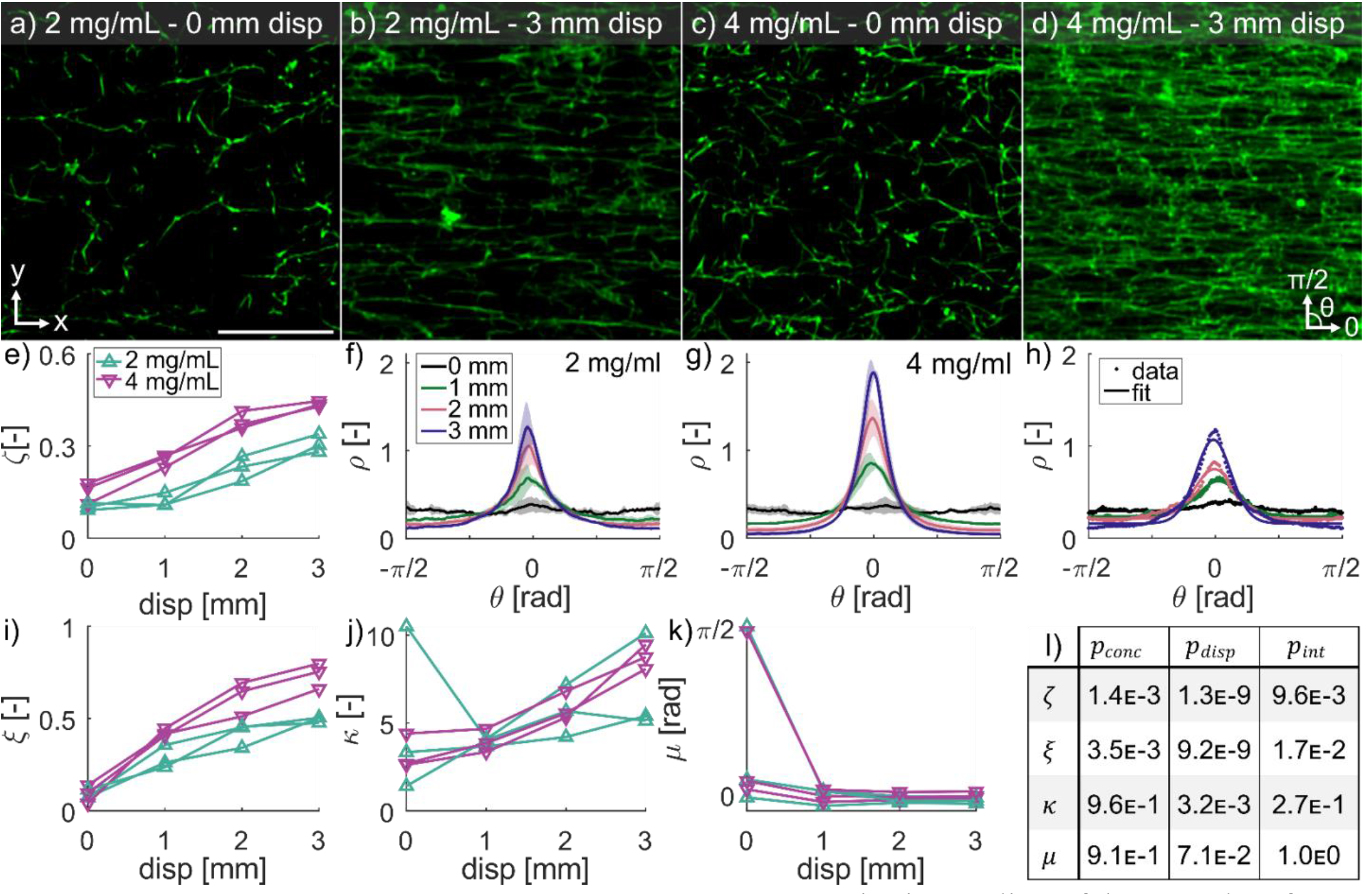
Fibrin aligns in the direction of deformation. Representative image slices of the *x* – *y* plane for a 2 mg/mL fibrin gel at 0 mm (a) and 3 mm (b) deformation, and a 4 mg/mL gel at 0 mm (c) and 3 mm (d) deformation in the *x*-direction (scale bar is 10 μm). Volume fraction of fibrin *ζ* was higher in 4 mg/mL gels and increased with increasing deformation (e). Mean +/− standard deviation of orientation distributions for 2 mg/mL (f) and 4 mg/mL (g) gels at increasing deformations showed strong realignment of fibers in the direction of deformation. Representative orientation distribution function fits at different displacements (h) show that the distribution function accurately captures the orientation distribution shape at all levels of displacement. The isotropic weighting parameter *ξ* increased with deformation (i) as the distributions became more anisotropic, and Von-Mises fiber alignment strength factor *κ* increased with deformation (j) as the fibers became more aligned in the x-direction. The direction of fiber alignment *μ* started near zero (x-direction) for most samples and quickly approached zero as the gels were deformed (k). The statistical analysis for concentration and displacement dependence, and the interaction between concentration and displacement for *ζ, ξ, κ*, and *μ* (l).

Estimated distribution parameters significantly varied as a function of both fibrinogen concentration and displacement. The isotropic weighting parameter *ξ* increased significantly with displacement (p=2.9E-9) (Fig. 4i), signifying an increase in anisotropy and differed depending on fibrinogen concentration (p=3.5E-3). The fiber alignment factor κ increased significantly with displacement (p=3.2E-3; Fig. 4j), indicating a more concentrated distribution in the direction of deformation, but was not affected by fibrinogen concentration (p=0.96). Mean fiber direction, μ, was not significantly affected by either displacement (p=0.071) or fibrinogen concentration (p=0.912; Fig. 4k). The fiber orientation was not affine (Fig. S1). The reorientation model, Eq.(8), was able to capture this non-affine deformation with the parameter Γ in the range 1.4 - 2.56. Individual fits of Γ for each gel are reported in Table S1 in the Supplement.

### 3.3 Stretch-dependent macroscale geometry

The 3D macroscale geometry of fibrin gels was measured between photobleached markers to evaluate the bulk deformation in response to applied stretch (Fig. 5a-c). The volume and cross-section of the region decreased with increasing stretch, by 92% and 88%, respectively (Fig. 5d,e). After 1 hour relaxation, significant decreases in cross-sectional area (p=0.040) and volume (p=0.018) were found; however, these were negligible in magnitude with decreases of approximately 1.1% and 1.5%, respectively (Fig. S2).

**Figure 5.**
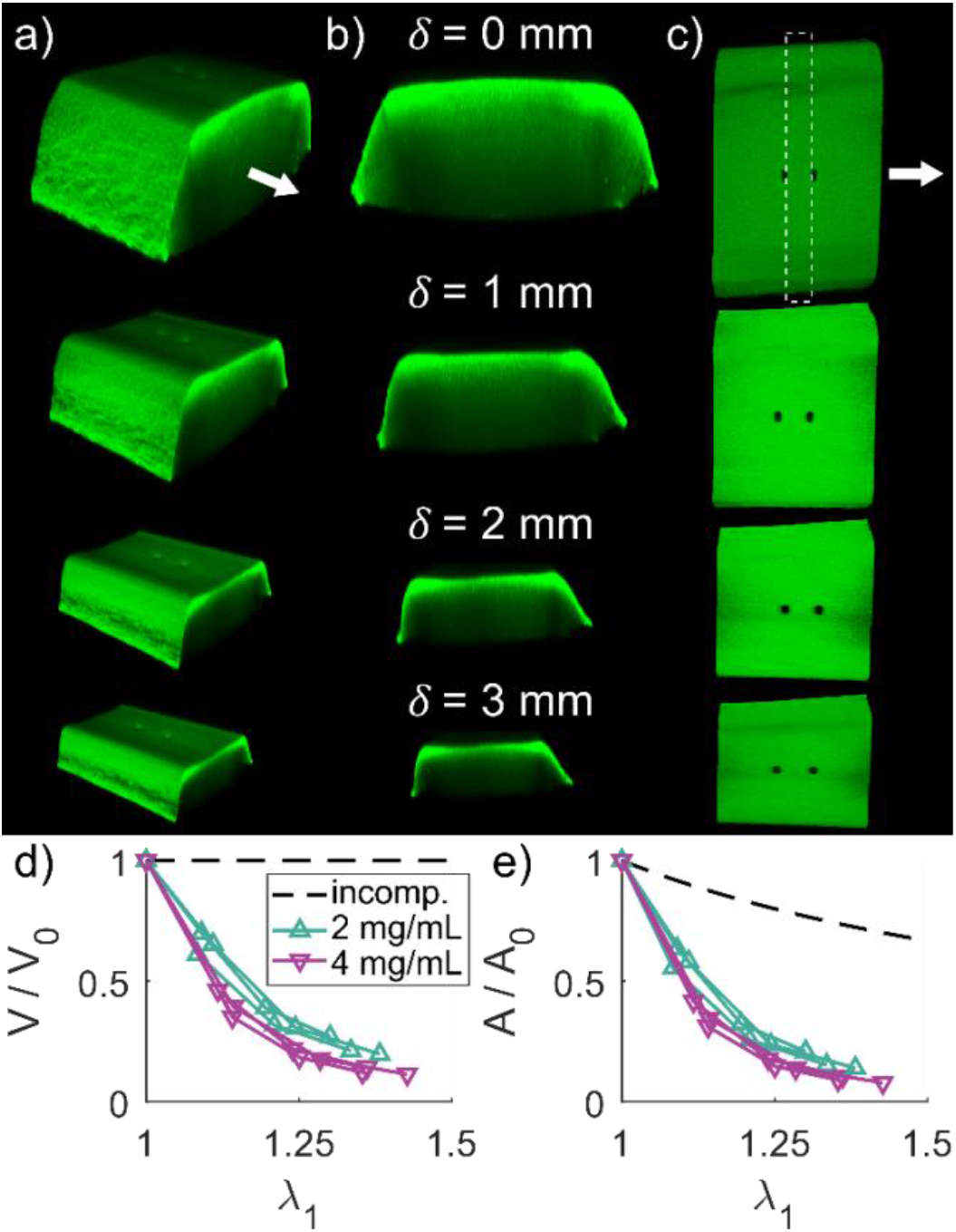
Fibrin gels substantially decreased in volume with deformation. Macroscale image stacks were taken at increasing levels of deformation to quantify changes in gel geometry. Isometric (a), cross-sectional (b) and top-down (c) views of a representative fibrin gel are shown for deformations of 0, 1, 2, and 3 mm. Stretch was defined by measuring the distance between photobleached markers as shown in (c, top). Volume was defined between the photobleached markers, and cross-sectional area was defined by all image stack slices through the length of the region between photobleached markers. Both relative volume (d) and relative cross-sectional area (e) decreased with increasing stretch for both 2 and 4 mg/mL gels. The dashed lines represent expected relative volume and cross-sectional area for an incompressible material.

### 3.4 Mechanical properties of fibrin gels

To quantify the viscoelastic response, fibrin gels underwent stress-relaxation tests. Stress decreased gradually over the 1 hour relaxation periods for both 2 and 4 mg/mL fibrin gels (Fig. 6a). The constitutive model Eqs. (2–7) captured the complete time-dependent behavior, including initial peak stress and relaxation response. To further characterize the viscoelastic response, initial and equilibrium stress values were isolated from the complete stress-relaxation data (Fig. 6b,c) and linear modulus fits were performed (Fig. 6d). The initial modulus and equilibrium modulus for the 2 mg/mL were 12.56 ± 7.50 kPa and 8.84 ± 4.79 kPa respectively, while for the 4 mg/mL the initial modulus was 31.83 ± 3.53 kPa and the equilibrium modulus was 23.12 ± 2.87 kPa. Thus, the 4 mg/mL gels were significantly stiffer than 2 mg/mL gels when considering both initial (p=0.0093) and equilibrium (p=0.0071) modulus.

**Figure 6.**
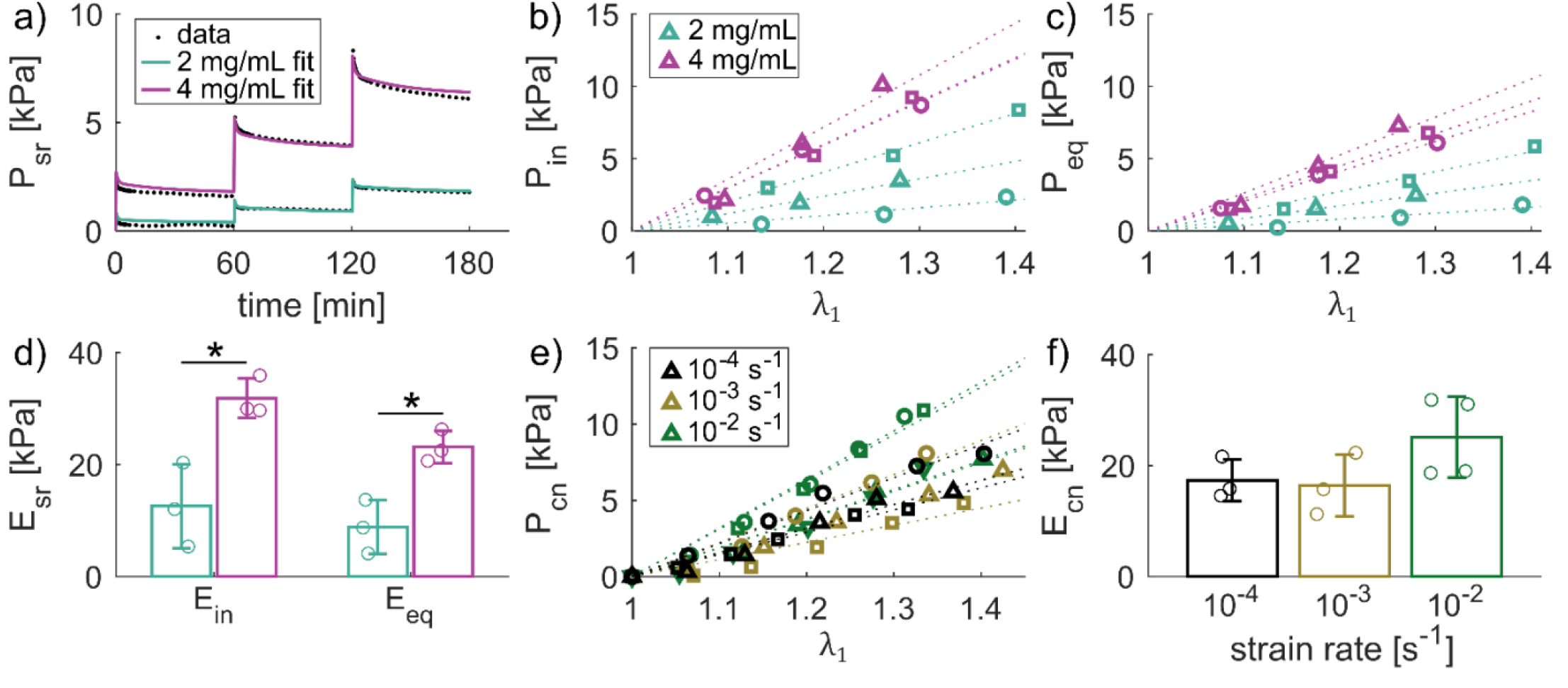
Fibrin gel tangential modulus increases with concentration. Fibrin gels showed viscoelastic behavior over the course of 1 hour relaxation steps, as seen by the representative 2 and 4 mg/mL gel stress-time responses (a). Initial stress response (b) and equilibrium stress (c) increased for each deformation step, with higher stresses in higher concentration gels. Tangential moduli fit to initial and equilibrium stress (dotted lines in b-c) were significantly higher for 4 mg/mL compared to 2 mg/mL gels. A subset of 2 mg/mL gels were deformed at different constant strain rates to similar levels of stretch (e). Tangential moduli were fit to the continuous stress-stretch data (dotted lines). Strain rate dependent modulus tended to increase with increasing strain rate but was not significant (f).

Tensile testing at different strain rates was performed to evaluate strain-rate dependence of the fibrin gel mechanical response (Fig. 6e). Although the stress-relaxation experiments exhibited a clear viscoelastic response, the tangent moduli from the strain-rate dependent tests were not significantly affected by strain rate (p=0.17; Fig. 6f).

### 3.5 Constitutive modeling

As described in ***2. Materials and Methods***, volume fraction (*ζ*), orientation distribution parameters (*ξ* and *κ*) and the reorientation parameter (Γ) were fit to the experimental results in Figs. 3 and 4 and are reported in Table S1. Then, using the data from Fig. 6, the rest of the parameters in Eqs. (2–7) were fit; *k*_0_, *k*_1_, *k*_2_, were determined to match the equilibrium stress data (Fig. S3a), and *β*_1_, *β*_2_, *τ*_1_, *τ*_2_ were optimized to match the viscoelastic response (Fig. S3b). The mean matrix and fiber moduli, *k*_0_ and *k*_1_ in Eqs. (2, 6), were 1352.73 Pa and 895.09 Pa, respectively, for the 2 mg/mL gels. For the 4mg/mL gels, the parameters *k*_0_, *k*_1_ had mean moduli of 3514.59Pa and 3455.67Pa, respectively. In line with the comparisons based on the tangent modulus, the fiber moduli parameters for the 4mg/mL gels were higher than those for the 2mg/mL gels. The nonlinearity of stress-stretch response was captured by the parameter *k*_2_, which was similar for the 2 mg/mL and 4 mg/mL gels, with means of 2.43 and 2.54, respectively. The mean equilibrium responses for the 2 mg/mL and 4 mg/mL gels are shown in Fig. S3c.

The stress relaxation response was accurately captured by the model by optimizing the parameters *β*_1_, *β*_2_, *τ*_1_, *τ*_2_ (Fig. 6a and Fig. S3b). Larger errors in the fit occurred at lower strains, possibly due to slack of the gel or boundary effects, with excellent accuracy for the second and third stretch steps. Fibrin gels showed two independent time scales of viscosity for both fibrinogen concentrations: a short-term relaxation time *τ*_1_ (51.7 s – 73.5 s for 2 mg/mL and 47.8 s – 66.5 s for 4 mg/mL gels) and a long-term relaxation time *τ*_2_ (1587.6 s – 1642.6 s for 2 mg/mL and 1664.3 s – 1702.2 s for 4 mg/mL gels). The average viscoelastic response is shown in Fig. S3d.

Average properties were then used in the finite element model (Fig. S3e) to verify that the simulations of the homogeneous gels matched the analytical solution of uniaxial tension (Eq. (S1)). In summary, the constitutive model and finite element implementation captured the mesoscale fiber organization, its non-affine deformation during tensile stretch, and the viscoelastic stress-strain response of homogeneous fibrin gels.

### 3.6 Testing and finite element analysis of heterogeneous fibrin gels

Two heterogeneous gels were fabricated as described in *Section 2.5*, and the corresponding finite element models were created following the methods in *Section 2.8*. For the 2 mg/mL gel with a 4 mg/mL inclusion (Fig. 7a), a non-uniform grid was photo-bleached around the inclusion and manually tracked as the gel was stretched by applying a 3000 μm displacement (Fig. 7b). The corresponding Green-Lagrange strain component in the direction of applied tension (*E_xx_*), showed larger strains in the 2 mg/mL region compared to the 4 mg/mL inclusion. In addition to tracking the deformation of the grid, the displacement of two photo-bleached markers near the PE blocks was also measured. The corresponding finite element model was created to match the geometry of the gel in the experiment (Fig. 7c). Average material parameters were used for the 2 mg/mL and 4 mg/mL subdomains, and the gel was stretched according to the displacement of the photo-bleached markers. The corresponding Green-Lagrange strain contour in the finite element simulation also showed larger strains in the 2 mg/mL region compared to the 4 mg/mL inclusion (Fig. 7d). The strain distributions in a gel made out of 4 mg/mL fibrin with a 2 mg/mL inclusion were more uniform in the experiments, with moderately larger strains in the 2 mg/mL inclusion compared to the surrounding 4 mg/mL material (Fig. 7e,f). The corresponding finite element model showed larger strain differences between the two subdomains, with the 2 mg/mL inclusion also undergoing larger deformation compared to the 4 mg/mL region (Fig. 7g,h).

**Figure 7.**
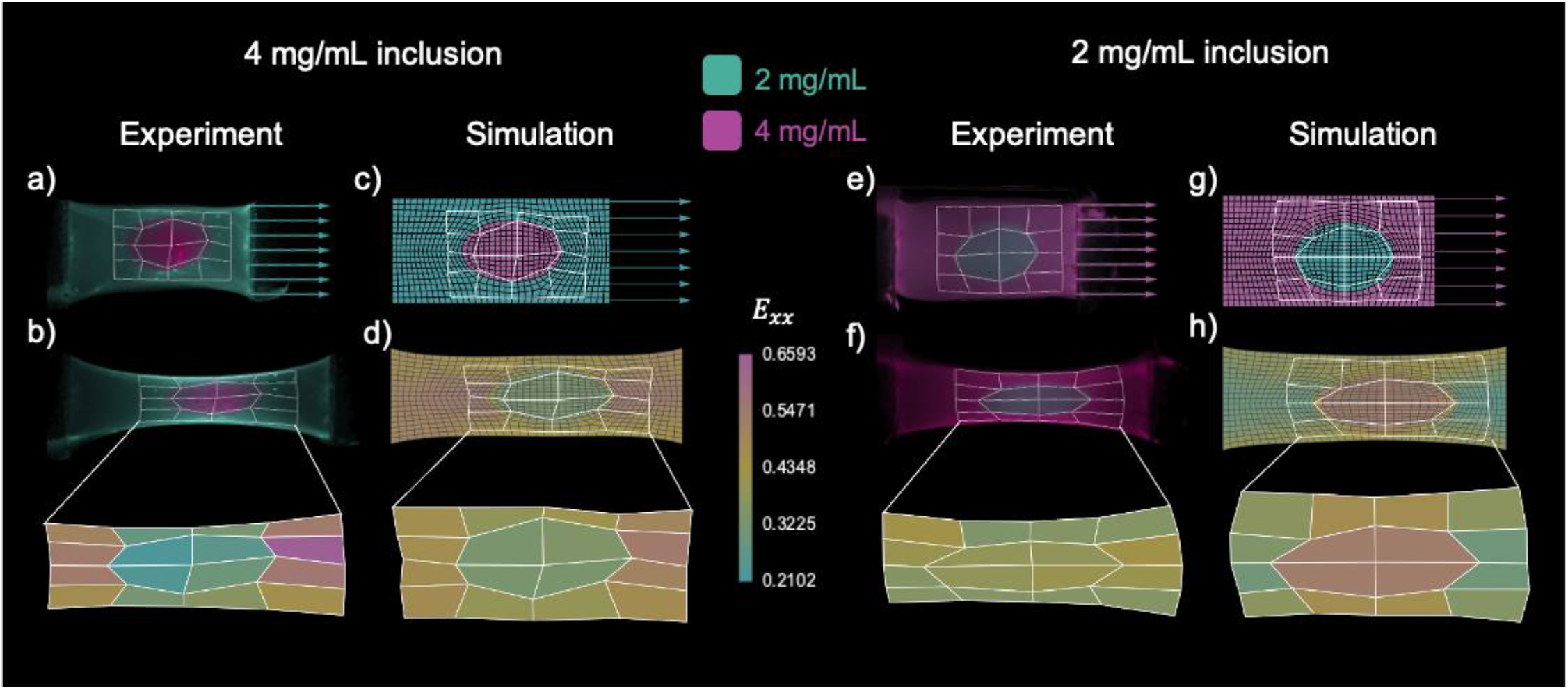
Computational models predict strain fields of heterogenous fibrin gels. A 2 mg/mL gel with a 4 mg/mL elliptical inclusion was stretched by applying a 3000 μm displacement on one end, and its deformation was tracked based on a photo-bleached grid (a). Unidirectional extension of the heterogeneous gel led to non-uniform strain *E_xx_*, with larger strains in the 2 mg/mL region with respect to the 4 mg/mL inclusion (b). A corresponding finite element model was created to replicate the geometry of the gel in the experiment, including the photo-bleached grid (c). Unidirectional extension of the finite element model using the average material parameters for the 2 mg/mL and 4 mg/mL homogeneous gels also showed heterogeneous strains, with larger strains in the 2 mg/mL domain compared to the inclusion. A 4 mg/mL gel with a 2 mg/mL inclusion was also constructed (e), and its strain measured by tracking a phot-bleached grid of quadrilaterals. The gel with the 2 mg/mL inclusion showed more homogeneous strain transfer, with moderately larger strains in the inclusion with respect to the surrounding domain. A finite element simulation matching the gel geometry was run for the gel with the 2 mg/mL inclusion (g). Finite element simulations showed greater variation in strain across domains compared to experiments, but showed larger strains in the 2mg/mL region, similar to experiments (h).

## 4 Discussion

Understanding the multiscale mechanical properties of fibrin matrices is an important step towards understanding the effects of loading and fiber remodeling during the wound healing process. Our study complements previous efforts on characterizing homogeneous fibrin gel mechanics; however, blood clots and dermal wounds are in a heterogeneous biomechanical environment, and the effect of domains of different material properties on the resulting mechanical response remains poorly understood. To address this gap, we conducted first a thorough characterization of homogeneous fibrin gels of different concentrations. The experiments on homogeneous gels were enhanced by the development and fitting of a nonlinear hyper-viscoelastic constitutive model and its implementation into a custom finite element solver. We then investigated heterogeneous fibrin gels relevant to dermal wounds by considering gels with elliptical inclusions of a different fibrin concentration. This study revealed that homogeneous fibrin gels show viscoelastic behavior, undergo large deformation-dependent volume loss, and present non-affine realignment of fibers in response to deformation. Experiments and modeling of heterogeneous gels showed that these gels develop heterogeneous strain fields across the regions with different fibrin concentrations. These methods can be expanded to study a myriad of biological interfaces, fiber matrices, and cell seeded constructs.

We first determined if fiber orientation in homogeneous gels was axially symmetric around the *x*-direction (long axis of the gel and direction of subsequent deformation; Fig. 3a-d). Undeformed fibrin gels polymerized with a slight preferential fiber orientation in the *x*-direction, likely the result of fluid flow when the fibrin solution was pipetted into the molds, as direction of flow has been shown to influence fiber orientation during fibrin polymerization [38,39]. Nevertheless, distribution parameters obtained when the data was fit to Eq. (1) revealed no significant differences in orientation based on fibrinogen concentration or rotational plane of interest. Therefore, our protocol for fibrin gel fabrication produces axially symmetric gels that can be modeled with transversely isotropic constitutive models.

Gels with higher fibrinogen concentration (4 mg/mL vs 2 mg/mL) had noticeably denser fiber matrices, which was supported by the significant difference in volume fraction found between concentrations (Fig. 4e). Surprisingly, the 4 mg/mL gels were not twice as dense as 2 mg/mL gel but were instead only 1.5-fold as dense on average when undeformed. This may be a result of either the less-dense matrices allowing more fluid flow out of the matrix during polymerization, a coupling between fiber network mechanics and equilibrium swelling, or a limitation of objective resolution. Additionally, the modulus was 2.6-fold higher for 4 mg/mL compared to 2 mg/mL gels (Fig 6d), indicating a nonlinear relationship between fibrinogen concentration, network structure, and gel mechanical response. Fibers in gels at both concentrations became aligned in the direction of deformation (Fig. 4a-d) that, when fit to Eq. (1), manifested as significant increases in the anisotropy parameters *ξ* and *κ* with displacement. While reorientation of fibers in the direction of deformation was expected, it was important to understand whether this reorientation was affine or not in order to accurately model the mechanical response of these biopolymer gels. Previous studies showed synthetic hydrogels and bio-polymer gels such as fibrin and collagen gels undergo non-affine reorientation [23], and modeling of discrete fiber networks suggests non-affine behavior [40]. Therefore, we tested whether the reorientation was affine by comparing predictions between an affine reorientation model (Eq. S3) and experimental data, and found that the affine deformation assumption was unable to capture the data (Fig. S1). We adopted a reorientation micromechanics model from [36] to describe the behavior observed in our experiments, which accurately captured the non-affine reorientation distribution with a single parameter (Fig. S1). Together, the initial fiber network density, non-affine reorientation during tensile stretching, and equilibrium stress-stretch response were fit accurately with a nonlinear multi-scale constitutive model. Importantly, our modeling framework bypassed non-affine discrete fiber network models that are computationally intensive [41], and instead used a micromechanics approach that was efficient and allowed for a custom finite element implementation (Fig S3e).

Stress-relaxation tests confirmed that fibrin gels behaved viscoelastically, with significant drop in the stress over one hour. The inferred time constants for the relaxation response agreed with previous work on fibrin gels, showing a rapid response for both fibrinogen concentrations [42–44], as well as a longer time-scale of relaxation [44]. The short-term time scale ranged from 47.8 s to 73.7 s, and the long-term time scale ranged from 1587.6 s to 1702.2 s. In contrast to the stress-relaxation response, strain-rate did not have a significant effect on modulus, though it trended to increase with increasing strain rate, consistent with the literature [44]. The hyper-viscoelastic model was able to accurately capture the overall stress-relaxation response of the gels (Figs. 6a, S3), and could easily be implemented by the custom finite element solver (Fig. S3e).

As fibrin gels were stretched, the significant decrease in volume of the region between photobleached markers revealed these gels were not incompressible (Fig. 5). The change in volume may be a result of substantial fluid loss between fibers or compression of the individual fibers. The increase in volume fraction of fibrin supports the idea that fluid was indeed lost, which was previously observed [4], and attributed to either fiber buckling normal to the direction of deformation, or that protein unfolding increased exposure to hydrophobic groups that expelled water and bundled stretched fibers. The mechanisms behind the compressible response were not accounted for in the modeling framework and remains a task of our ongoing work.

Having characterized homogeneous gels, we proceeded to test fibrin gels of a given concentration with an inclusion made out of a different concentration of fibrin (Fig. 7). The heterogeneous gels formed stable interfaces between domains with different fibrin concentrations, resulting in the continuous deformation of the gel under tension. In other words, there was no evident fracture or debonding between the inclusion and the surrounding material. The strain distributions observed experimentally for the heterogeneous gels were compared with finite element simulations informed only by the homogeneous gel data. The experiment and simulation both showed that a soft inclusion undergoes larger strains compared to the stiffer surrounding material and, vice versa, that stiff inclusions deform less than the surrounding material. However, the variation in strain predicted by the finite element simulations was much larger than that observed in experiments. This was due, at least partially, to the fact that the inclusions did not occupy the entire depth of the gels, in contrast with the finite element simulations. Nevertheless, the model and experiments suggest that heterogeneous fibrin gels can be constructed to achieve corresponding heterogeneous strain distribution by means of inclusions. This type of heterogeneous gel design is not only needed to capture physiological scenarios, but also opens the possibility for the design of fibrin gels with controllable mesoscale strain concentrations for scaffold design.

## 5 Conclusions

Measuring and modeling the mechanics of fibrin matrices is important for building a foundation of knowledge of the wound healing process and furthering the development of regenerative scaffolds. The characteristics of the fibrin networks we uncovered in this work (*e.g*. concentration dependent mechanics, non-affine fiber reorientation, and deformation induced volume loss and volume fraction increase) provide useful targets for constitutive models to capture. Our proposed constitutive model captured the mechanical response of the fibrin gels while predicting the strain fields of heterogeneous concentration fibrin gel constructs. The model and mechanical characterization methods presented in this study can be utilized and built upon to study cell seeded matrices, heterogeneous matrices that more closely mimic the wounds healing environment (*e.g*. collagen gels with fibrin gel inclusions), and more complex matrix organizations resulting from the advancement of 3D printing and microfluidic-based polymerization methods. Increased knowledge of the underlying properties of polymerized biological gel matrices through evolution and refinement of methods of gel construction, mechanical analysis, and constitutive modeling will pave the way for development of new innovative treatment strategies in the wound healing space.

## Acknowledgements

This work was supported by NSF 1911346-CMMI to S.C. and A.B.T. and NIH DP2 AT009833 to S.C. The authors would like to thank Dr. Robert MacCurdy for the use of the Stratasys J750 3D printer and Trotec Speedy 360 Laser Cutter, Brandon Hayes and Lawrence Smith for assistance with 3D printing fibrin gel molds, and members of the Calve lab, including Dr. Callan Luetkemeyer, for helpful discussions.

## Data Availability

Data presented in this study is available upon request from the corresponding authors.

**Appendices (optional)**

## Nomenclature

| Symbol | Description |
| --- | --- |
| $\rho$ | Probability density function |
| $\theta$ | Fiber orientation |
| $\xi$ | Isotropic weighting parameter |
| $\kappa$ | Fiber alignment strength factor |
| $\mu$ | Preferred fiber orientation |
| $I_0$ | Modified Bessel function of the first kind of order zero |
| $\lambda$ | Stretch |
| $\zeta$ | Volume fraction of fibrin |
| $\delta$ | Applied unidirectional deformation |
| $V$ | Volume between photobleached fibrin gel markers |
| $V_0$ | Initial volume between photobleached fibrin gel markers |
| $A$ | Mean cross-sectional area between photobleached fibrin gel markers |
| $A_0$ | Initial mean cross-sectional area volume between photobleached fibrin gel markers |
| $P$ | First Piola-Kirchhoff stress |
| $P_{sr}$ | First Piola-Kirchhoff stress for stress-relaxation |
| $P_{cn}$ | First Piola-Kirchhoff stress for constant strain-rate deformation |
| $E_{sr}$ | Tangent modulus for stress-relaxation |
| $E_{in}$ | Initial stress tangent modulus for stress-relaxation |
| $E_{eq}$ | Final stress tangent modulus for stress-relaxation |
| $E_{cn}$ | Tangent modulus for constant strain-rate deformation |
| $E_{11}$ | Green-Lagrange strain in the direction of applied deformation |
| $k_1$ | Fiber modulus in Eq. (2) |
| $k_2$ | Fiber nonlinearity in Eq. (2) |
| $\beta_1, \beta_2$ | Scaling coefficients for the viscoelastic energy in Eq. (2) |
| $\tau_1, \tau_2$ | Time scales of relaxation for the viscoelastic response in Eq. (2) |
| $k_0$ | Modulus of the ECM *ground substance* in Eq. (6) |
| $\chi$ | Reorientation function defined in Eq. (8) |
| $\theta_c$ | Direction of principal stretch |
| $\lambda_c$ | Magnitude of the principal stretch |
| $\Gamma$ | Parameter controlling the non-affine reorientation |

## Supplementary material

### First Piola Kirchhoff stress

The main text introduced energy expressions Eqs. (2–7) which fully define the mechanical response of the gels. For data fitting, calculation of the first Piola Kirchhoff stress, ***P***, is necessary to compare model predictions to experiments. The expression for the first component of ***P*** under uniaxial deformation based on Eqs. (2–7) is

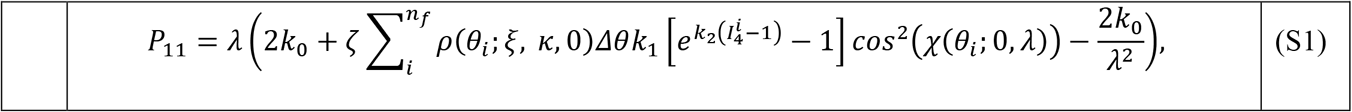

Where *λ* is the applied uniaxial stretch. The assumption of uniaxial stretch has been used in the evaluation of the reorientation function, *χ*(*θ_i_*; 0, *λ*), by setting *θ_c_* = 0, *λ_c_* = *λ*. The uniaxial assumption is also employed to obtain the stretch for each fiber based on

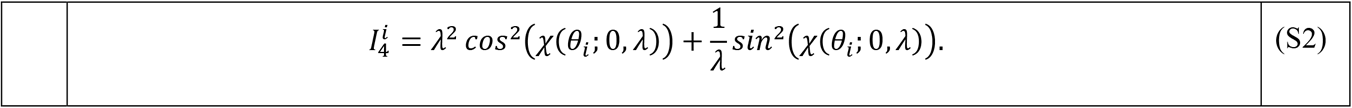

### Affine versus non-affine reorientation

Affine reorientation is described by the following change in angle with deformation

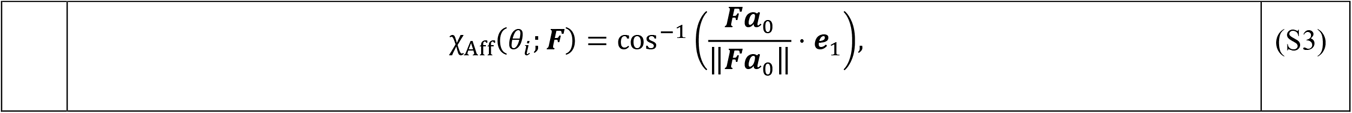

where ***a***_0_ = [cos(*θ_i_*), sin(*θ_i_*)], is the initial fiber orientation, ***F*** is the deformation gradient, and ***e**_A_* = [1,0] is the unit vector in the direction of uniaxial loading. Non-affine reorientation is described with Eq.(8) in the main text.

**Fig S1.**
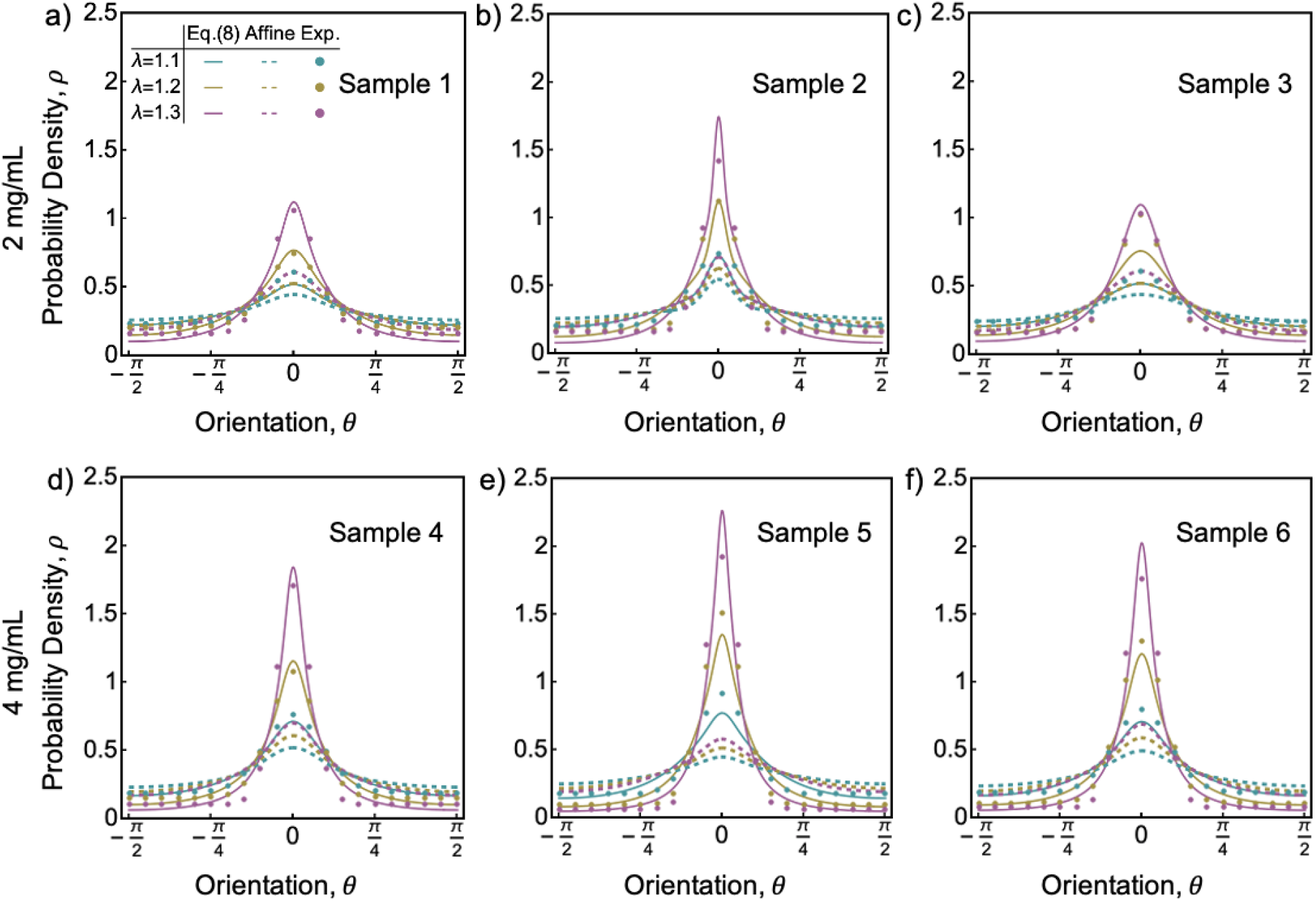
Fibrin fiber distribution changes non-affinely with increasing tensile stretch. Modeling the reorientation as affine (Eq. (S3)), is unable to capture the experimental results for either the 2 mg/mL (a) and 4 mg/mL (b). In contrast, Eq. (8) was able to account for non-affine reorientation, accurately matching the experimental data. Due to non-affine deformation, fibers become preferentially oriented around the direction of applied stretch, *θ_c_* = 0, as the stretch *λ* increases.

**Figure S2.**
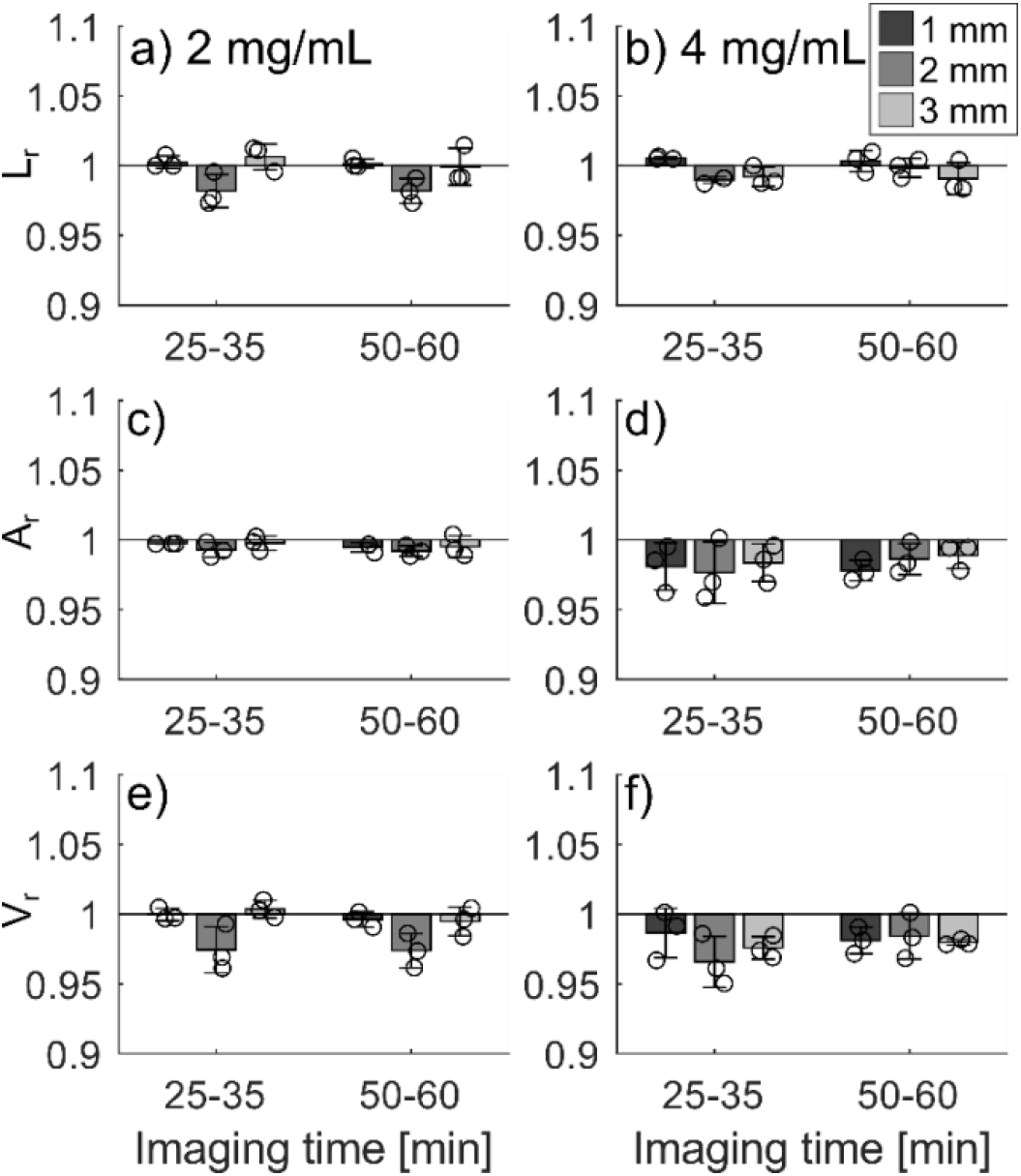
There were negligible time-dependent changes in geometry of fibrin gels. Full thickness images stacks of the fibrin gels were acquired at 0-5, 25-35, and 50-60 minutes after each deformation. Length between photobleached markers (a,b), mean cross-sectional area (c,d), and volume (e,f) were analyzed at the different timepoints and normalized to values obtained at the initial 0-5 minute timepoints. Statistical analysis for each measure consisted of a 3-way ANOVA with concentration, displacement, and imaging time as sources of variation, and a Tukey-Kramer test for post-hoc analysis. Significance was found between initial (0-5 minutes) and final (50-60 minutes) imaging times for cross-sectional area (p=0.040) and volume (p=0.018). As expected, length, cross-sectional area, and volume were significantly different between all displacements (p<0.01). Additionally, significance was found between concentrations for sample length (p=0.049). While significance was found between initial and final time points for area and volume, the magnitude of the difference was small enough on average (~1.3%) that we deemed the difference negligible for modeling purposes.

**Table S1.** Constitutive model parameters. Mesoscale parameters were obtained from experiments characterizing the fiber network structure and deformation during unidirectional tensile stretch: *ζ_f_*, volume fraction; *ξ*, isotropic orientation distribution weight; *κ*, concentration parameter for the von-Mises orientation distribution; Γ, reorientation parameter to capture the non-affine alignment of fibers in the direction of the principal stretch. From the tensile and stress relaxation tests, the following parameters associated with the equilibrium stress response were used: *k*_0_, shear modulus of the neo-Hookean ground substance; *k*_1_, shear modulus of individual fibers; *k*_2_, a dimensionless parameter that captures nonlinearity in the response. The parameters associated with the viscoelastic dissipation response are: *β*_1_, *β*_2_, dimensionless factors that describe the contribution of the viscous branches to the total stress as proportional to the equilibrium branches; *τ*_1_, *τ*_2_, time constants of the stress relaxation response. Length-scale parameters were used as regularization to balance between individual sample fit and a single fit to all data and reflect the expected range for the parameters based on a first optimization run without correlation terms.

| | $k_0$ [Pa] | $k_1$ [Pa] | $k_2$ [-] | $\zeta_f$ | $\xi$ | $\kappa$ | $\Gamma$ | $\beta_1$ | $\tau_1$ [s] | $\beta_2$ | $\tau_2$ [s] |
| --- | --- | --- | --- | --- | --- | --- | --- | --- | --- | --- | --- |
| Sample 1 | 2124.16 | 1017.71 | 2.47674 | 0.2031 | 0.3915 | 2.6344 | 1.56119 | 8.84 | 63.5 | 4.75 | 1626.4 |
| Sample 2 | 367.58 | 856.07 | 2.59243 | 0.1342 | 0.0498 | 6.3612 | 2.14685 | 4.59 | 51.7 | 2.42 | 1642.6 |
| Sample 3 | 1566.45 | 811.49 | 2.20447 | 0.1550 | 0.4138 | 2.9137 | 1.3999 | 7.83 | 73.7 | 5.87 | 1587.6 |
| Average | 1352.73 | 895.09 | 2.4245 | 0.1641 | 0.2850 | 3.9698 | 1.70265 | 7.09 | 63.0 | 4.35 | 1618.9 |
| Length-scale $s$ | 3000 | 100 | 0.5 | -- | -- | -- | -- | 10 | 50 | 10 | 200 |
| Sample 5 | 3143.56 | 3417.75 | 2.53692 | 0.1847 | 0.4443 | 8.4585 | 1.98479 | 4.25 | 57.6 | 1.66 | 1689.7 |
| Sample 6 | 3761.51 | 3405.98 | 1.25 | 0.1982 | 0.4659 | 4.0164 | 2.56463 | 6.37 | 47.8 | 5.76 | 1664.3 |
| Sample 7 | 3638.69 | 3543.28 | 3.81909 | 0.1485 | 0.1174 | 3.0523 | 1.88596 | 6.79 | 66.5 | 3.17 | 1702.2 |
| Average | 3514.59 | 3455.67 | 2.5353 | 0.1771 | 0.3425 | 5.1757 | 2.14513 | 5.80 | 57.3 | 3.53 | 1685.4 |
| Length-scale $s$ | 3000 | 220 | 2 | -- | -- | -- | -- | 10 | 50 | 10 | 200 |

**Figure S3:**
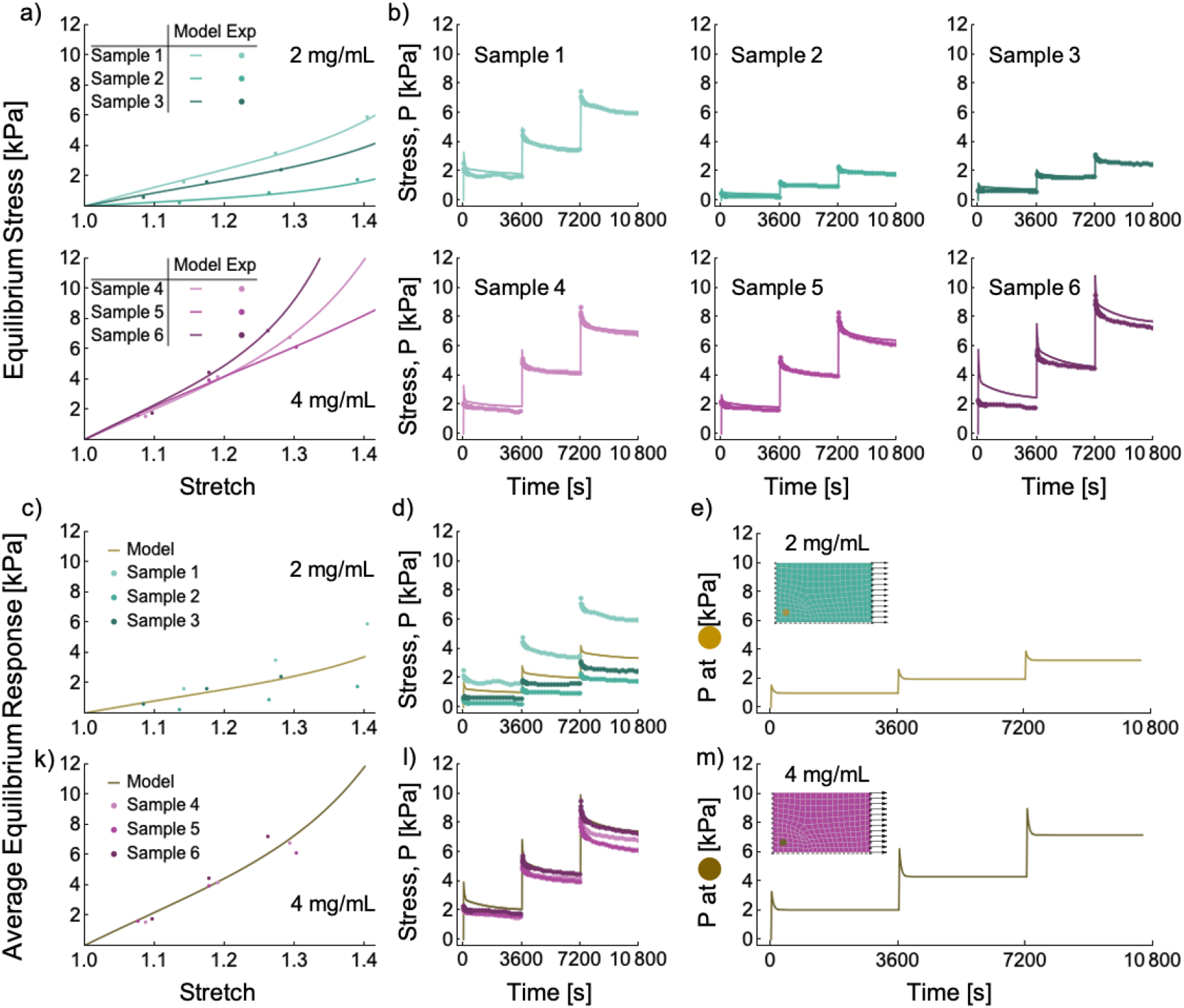
Fit of constitutive model to experimental data and finite element verification. The individual parameter fits from Table S1 were used to model the relaxed stress response for 2 mg/mL and 4 mg/mL gels (a), as well as the stress relaxation response (b). The average parameters captured the relaxed stress and viscoelastic relaxation trends for the 2mg/mL and for 4mg/mL (c, d) compared to experiments. Homogeneous fibrin gels were modeled with a custom C++ finite element implementation using the average parameters and boundary conditions set to mimic the tensile tests from experiments and verify the model implementation (e).

## References

[1] P.A. Janmey, J.P. Winer, J.W. Weisel, Fibrin gels and their clinical and bioengineering applications, J R Soc Interface. 6 (2009) 1–10. https://doi.org/10.1098/rsif.2008.0327.

[2] R.I. Litvinov, J.W. Weisel, Fibrin mechanical properties and their structural origins, Matrix Biology. 60–61 (2017) 110–123. https://doi.org/10.1016/j.matbio.2016.08.003.

[3] G. Dietler, W. Känzig, A. Haeberli, P.W. Straub, Temperature dependence of fibrin polymerization: a light scattering study, Biochemistry. 24 (1985) 6701–6706. https://doi.org/10.1021/BI00344A060.

[4] A.E.X. Brown, R.I. Litvinov, D.E. Discher, P.K. Purohit, J.W. Weisel, Multiscale mechanics of fibrin polymer: Gel stretching with protein unfolding and loss of water, Science (1979). 325 (2009) 741–744. https://doi.org/10.1126/science.1172484.

[5] C.L. Gilchrist, D.S. Ruch, D. Little, F. Guilak, Micro-scale and meso-scale architectural cues cooperate and compete to direct aligned tissue formation, (2014). https://doi.org/10.1016/j.biomaterials.2014.08.047.

[6] A.E.X. Brown, R.I. Litvinov, D.E. Discher, J.W. Weisel, Forced unfolding of coiled-coils in fibrinogen by single-molecule AFM, Biophys J. 92 (2007). https://doi.org/10.1529/biophysj.106.101261.

[7] W. Li, J. Sigley, S.R. Baker, C.C. Helms, M.T. Kinney, M. Pieters, P.H. Brubaker, R. Cubcciotti, M. Guthold, Nonuniform Internal Structure of Fibrin Fibers: Protein Density and Bond Density Strongly Decrease with Increasing Diameter, (2017). https://doi.org/10.1155/2017/6385628.

[8] S. Yesudasan, R.D. Averett, Multiscale network modeling of fibrin fibers and fibrin clots with protofibril binding mechanics, Polymers (Basel). 12 (2020). https://doi.org/10.3390/POLYM12061223.

[9] F. Maksudov, A. Daraei, A. Sesha, K.A. Marx, M. Guthold, V. Barsegov, Strength, deformability and toughness of uncrosslinked fibrin fibers from theoretical reconstruction of stress-strain curves, Acta Biomater. 136 (2021) 327–342. https://doi.org/10.1016/J.ACTBIO.2021.09.050.

[10] C.J.G. Abrego, L. Dedroog, O. Deschaume, J. Wellens, A. Vananroye, M.P. Lettinga, J. Patterson, C. Bartic, Multiscale Characterization of the Mechanical Properties of Fibrin and Polyethylene Glycol (PEG) Hydrogels for Tissue Engineering Applications, Macromol Chem Phys. 223 (2022). https://doi.org/10.1002/macp.202100366.

[11] J.W. Weisel, The mechanical properties of fibrin for basic scientists and clinicians, 112 (2004) 267–276. https://doi.org/10.1016/j.bpc.2004.07.029.

[12] E.A. Ryan, L.F. Mockros, J.W. Weisel, L. Lorand, Structural origins of fibrin clot rheology, Biophys J. 77 (1999) 2813–2826. https://doi.org/10.1016/S0006-3495(99)77113-4.

[13] W.W. Roberts, O. Kramer, R.W. Rosser, F.H.M. Nestler, J.D. Ferry, Rheology of fibrin clots. I.: Dynamic viscoelastic properties and fluid permeation, Biophys Chem. 1 (1974) 152–160. https://doi.org/10.1016/0301-4622(74)80002-5.

[14] S. Nam, K.H. Hu, M.J. Butte, O. Chaudhuri, Strain-enhanced stress relaxation impacts nonlinear elasticity in collagen gels, Proc Natl Acad Sci U S A. 113 (2016) 5492–5497. https://doi.org/10.1073/PNAS.1523906113/SUPPL_FILE/PNAS.201523906SI.PDF.

[15] H. Wagreich, I. Tarlov, Studies on the strength of fibrinogen-thrombin clots, Arch Biochem. 7 (1945) 345–352.

[16] W.W. Roberts, L. Lorand, L.F. Mockros, Viscoelastic properties of fibrin clots, Biorheology. 10 (1973) 29–42. https://doi.org/10.3233/BIR-1973-10105.

[17] E. Fukada, L. Dintenfass, The clotting of blood and fibrinogen-thrombin systems as studied by two dynamic instruments with large and small amplitudes, Biorheology. 8 (1971) 149–155. https://doi.org/10.3233/BIR-1971-83-405.

[18] M. Kaibara, E. Fukada, Dynamic viscoelastic study for the structure of fibrin networks in the clots of blood and plasma, Biorheology. 6 (1970) 329–339. https://doi.org/10.3233/BIR-1970-6407.

[19] J.D. Ferry, M. Miller, S. Shulman, The conversion of fibrinogen to fibrin. VII. Rigidity and stress relaxation of fibrin clots; effect of calcium, Arch Biochem Biophys. 34 (1951) 424–436. https://doi.org/10.1016/0003-9861(51)90021-5.

[20] K.B. Neeves, D.A.R. Illing, S.L. Diamond, Thrombin flux and wall shear rate regulate fibrin fiber deposition state during polymerization under flow, Biophys J. 98 (2010) 1344–1352. https://doi.org/10.1016/j.bpj.2009.12.4275.

[21] P. Whittaker, K. Przyklenk, Fibrin architecture in clots: a quantitative polarized light microscopy analysis, Blood Cells Mol Dis. 42 (2009) 51–56. https://doi.org/10.1016/J.BCMD.2008.10.014.

[22] D.G. Norton, N.K. Fan, M.J. Goudie, H. Handa, M.O. Platt, R.D. Averett, Computational imaging analysis of glycated fibrin gels reveals aggregated and anisotropic structures, J Biomed Mater Res A. 105 (2017) 2191–2198. https://doi.org/10.1002/JBM.A.36074.

[23] Q. Wen, A. Basu, P.A. Janmey, A.G. Yodh, Non-affine deformations in polymer hydrogels, Soft Matter. 8 (2012) 8039–8049. https://doi.org/10.1039/C2SM25364J.

[24] M. Aghvami, V.H. Barocas, E.A. Sander, Multiscale Mechanical Simulations of Cell Compacted Collagen Gels, J Biomech Eng. 135 (2013) 0710041. https://doi.org/10.1115/1.4024460.

[25] E.A. Sander, V.H. Barocas, R.T. Tranquillo, Initial fiber alignment pattern alters extracellular matrix synthesis in fibroblast-populated fibrin gel cruciforms and correlates with predicted tension, Ann Biomed Eng. 39 (2011) 714–729. https://doi.org/10.1007/s10439-010-0192-2.

[26] A.M. De Jesus, M. Aghvami, E.A. Sander, A Combined In Vitro Imaging and Multi-Scale Modeling System for Studying the Role of Cell Matrix Interactions in Cutaneous Wound Healing, PLoS One. 11 (2016). https://doi.org/10.1371/JOURNAL.PONE.0148254.

[27] A.S. Abhilash, B.M. Baker, B. Trappmann, C.S. Chen, V.B. Shenoy, Remodeling of Fibrous Extracellular Matrices by Contractile Cells: Predictions from Discrete Fiber Network Simulations, Biophysj. 107 (2014) 1829–1840. https://doi.org/10.1016/j.bpj.2014.08.029.

[28] E. Ban, H. Wang, J. Matthew Franklin, J.T. Liphardt, P.A. Janmey, V.B. Shenoy, Strong triaxial coupling and anomalous Poisson effect in collagen networks, Proc Natl Acad Sci U S A. 116 (2019) 6790–6799. https://doi.org/10.1073/pnas.1815659116.

[29] Y. Leng, V. Tac, S. Calve, A.B. Tepole, Predicting the mechanical properties of biopolymer gels using neural networks trained on discrete fiber network data, Comput Methods Appl Mech Eng. 387 (2021) 114160. https://doi.org/10.1016/J.CMA.2021.114160.

[30] Y. Guo, S. Calve, A.B. Tepole, Multiscale mechanobiology: Coupling models of adhesion kinetics and nonlinear tissue mechanics, Biophys J. 121 (2022) 525–539. https://doi.org/10.1016/J.BPJ.2022.01.012.

[31] A. Acuna, J.M. Jimenez, N. Deneke, S.M. Rothenberger, S. Libring, L. Solorio, V.L. Rayz, C.S. Davis, S. Calve, Design and validation of a modular micro-robotic system for the mechanical characterization of soft tissues, Acta Biomater. 134 (2021) 466–476. https://doi.org/10.1016/J.ACTBIO.2021.07.035.

[32] E.E. Morrill, A.N. Tulepbergenov, C.J. Stender, R. Lamichhane, R.J. Brown, T.J. Lujan, A validated software application to measure fiber organization in soft tissue, Biomech Model Mechanobiol. 15 (2016) 1467–1478. https://doi.org/10.1007/S10237-016-0776-3.

[33] C.L.M. Gouget, M.J. Girard, C.R. Ethier, A constrained von Mises distribution to describe fiber organization in thin soft tissues, Biomech Model Mechanobiol. 11 (2012) 475–482. https://doi.org/10.1007/S10237-011-0326-Y.

[34] G.A. Holzapfel, T.C. Gasser, R.W. Ogden, A New Constitutive Framework for Arterial Wall Mechanics and a Comparative Study of Material Models, Journal of Elasticity and the Physical Science of Solids 2000 61:1. 61 (2000) 1–48. https://doi.org/10.1023/A:1010835316564.

[35] H. Liu, G.A. Holzapfel, B.H. Skallerud, V. Prot, Anisotropic finite strain viscoelasticity: Constitutive modeling and finite element implementation, J Mech Phys Solids. 124 (2019) 172–188. https://doi.org/10.1016/J.JMPS.2018.09.014.

[36] A. Raina, C. Linder, A homogenization approach for nonwoven materials based on fiber undulations and reorientation, J Mech Phys Solids. 65 (2014) 12–34. https://doi.org/10.1016/J.JMPS.2013.12.011.

[37] P. Knysh, Y. Korkolis, Blackbox: A procedure for parallel optimization of expensive black-box functions, (2016). https://doi.org/10.48550/arxiv.1605.00998.

[38] R.I. Litvinov, J.W. Weisel, Fibrin mechanical properties and their structural origins, Matrix Biology. 60–61 (2017) 110–123. https://doi.org/10.1016/J.MATBIO.2016.08.003.

[39] R.A. Campbell, M.M. Aleman, L.D. Gray, M.R. Falvo, A.S. Wolberg, Flow profoundly influences fibrin network structure: Implications for fibrin formation and clot stability in haemostasis, Thromb Haemost. 104 (2010) 1281–1284. https://doi.org/10.1160/TH10-07-0442.

[40] P.L. Chandran, V.H. Barocas, Affine versus non-affine fibril kinematics in collagen networks: theoretical studies of network behavior, J Biomech Eng. 128 (2006) 259–270. https://doi.org/10.1115/1.2165699.

[41] A. Mauri, R. Hopf, A.E. Ehret, C.R. Picu, E. Mazza, A discrete network model to represent the deformation behavior of human amnion, J Mech Behav Biomed Mater. 58 (2016) 45–56. https://doi.org/10.1016/J.JMBBM.2015.11.009.

[42] K. Tashiro, Y. Shobayashi, I. Ota, A. Hotta, Finite element analysis of blood clots based on the nonlinear visco-hyperelastic model, (2021). https://doi.org/10.1016/j.bpj.2021.08.034.

[43] S. Johnson, R. McCarthy, M. Gilvarry, P.E. McHugh, J.P. McGarry, Investigating the Mechanical Behavior of Clot Analogues Through Experimental and Computational Analysis, Ann Biomed Eng. 49 (2021) 420–431. https://doi.org/10.1007/S10439-020-02570-5.

[44] G.P. Sugerman, S.H. Parekh, M.K. Rausch, Nonlinear, dissipative phenomena in whole blood clot mechanics, Soft Matter. 16 (2020) 9908–9916. https://doi.org/10.1039/D0SM01317J.

